# An 8 Dipole Transceive and 24 Loop Receive Array for Non-Human Primate Head Imaging at 10.5T

**DOI:** 10.1101/2020.08.27.270744

**Authors:** Russell L. Lagore, Steen Moeller, Jan Zimmermann, Lance DelaBarre, Jerahmie Radder, Andrea Grant, Kamil Ugurbil, Essa Yacoub, Noam Harel, Gregor Adriany

## Abstract

A 32-channel RF coil was developed for brain imaging of anesthetized non-human primates (Rhesus Macaque) at 10.5 tesla. The coil is composed of an 8-channel dipole transmit/receive array, close-fitting 16-channel loop receive array headcap, and 8-channel loop receive array lower insert.

The transceiver dipole array is composed of eight end-loaded dipole elements self-resonant at the 10.5 tesla proton Larmor frequency. These dipole elements were arranged on a plastic cylindrical former which was split in two to allow for convenient animal positioning. Nested into the bottom of the dipole array former is located an 8-channel loop receive array which contains 5×10 cm^2^ square loops arranged in two rows of four loops. Arranged in a close-fitting plastic headcap is located a high-density 16-channel loop receive array. This array is composed of 14 round loops 37 mm in diameter and two partially detachable, irregularly shaped loops that encircle the ears. Imaging experiments were performed on anesthetized non-human primates on a 10.5 tesla MRI system equipped with body gradients with a 60 cm open bore.

The coil enabled submillimeter (0.58 mm isotropic) high resolution anatomical and functional imaging as well as tractography of fasciculated axonal bundles. The combination of a close-fitting loop receive array and dipole transceiver array allowed for a higher channel count receiver and consequent higher SNR and parallel imaging gains. Parallel imaging performance supports high resolution fMRI and dMRI with a factor of three reduction in sampling. The transceive array elements during reception contributed approximately one quarter of SNR in the lower half of the brain which was farthest from the close-fitting headcap receive array.

## Introduction

In recent years, there has been a proliferation of ultra-high field (UHF) systems (defined as 7T and higher) for human imaging, primarily motivated by the fact that intrinsic gains in signal-to-noise ratio (SNR) and, in many cases, contrast-to-noise ratio (CNR) with increasing field strength will permit the acquisition of anatomical, functional and connectivity information with higher spatial and temporal resolutions. This in turn allows better characterization of the functional architectures and the underlying cognitive processes in the brain. Naturally, UHF systems capable of imaging humans have enormous appeal to the vast neuroscience community including the study of non-human primates (NHP).^1–4^ However, the ability to realize the gains that come with UHF is predicated on the availability of optimized RF coils.^5–9^ At clinical field strengths (1.5T and 3T), optimized RF coils tend to be receive-only array designs which utilize a large behind the bore liner RF body coil for excitation; whereas at UHF, body transmit RF coils are not typically used and thus local transmit and receive coil array designs are needed. Similar to human head imaging at UHF, a combinations of transmitter arrays with receiver arrays are desirable for whole head primate applications since they can address transmit inhomogeneity issues at UHF and optimal SNR simultaneously. Gilbert et al.^5^ followed this path and presented a compact 8 channel transmit, 24 channel receive array for 7T primate imaging within a head gradient coil designed for human head imaging. In previous work^10^ we described the development of a 16-channel decoupled transmission line transceiver with 6-channel loop receiver for imaging the brain of NHPs at 7T. This design proved successful due to the SNR benefits of a close-fitting loop receive-only array in combination with a “transceiver” array, constructed of stripline elements, that operated both for transmission and reception. This is an appealing concept since the unique field sensitivity profiles of the striplines contributed both to improved parallel imaging performance and supported whole head coverage. The transceiver concept that utilized the stripline array during transmission and reception (no active transmitter detuning), was achievable since the striplines can be positioned to be decoupled from the loop elements of the receive array.

In this paper, we present the development of an NHP coil for 10.5T (447 MHz), the highest magnetic field system that has recently become available for human or large animal imaging^11^. We extend the concept of using combined transceiver and receive only arrays in this case utilizing dipole transceivers combined with 8, 16 and 24 channel classical loop receivers. There are several benefits to using dipole elements at this high magnetic field compared to stipline elements. Advantages of dipole arrays have been demonstrated at 7T and 10.5T for imaging the human abdomen and head.^12,13^ Other works have demonstrated the advantages of a loop-dipole combination.^14,15^ In this work, we utilize an 8-channel transceiver dipole array together with an 8-channel and a 16-channel loop receive only arrays for an effective 8 channel transmit and 32 channel receive coil. A combination of transceiver dipoles with receive loop arrays increases the number of decoupled receiver channels and is expected to optimize penetration and SNR for obtaining high resolutions of deep and sub-cortical structures. The aim of the study was to build and evaluate this combination of transmit and receive arrays for NHP applications.

## Methods

A cylindrical coil former (19 cm ID), split into two pieces along the central axis, was designed in Blender 2.79 (Blender Foundation, Amsterdam, Netherlands) and fabricated with an F400 3D printer (Fusion3, Greensboro, NC, USA). This housing accommodates eight 18 cm long end-loaded dipole antennas and features integrated ear bars to allow for consistent positioning of NHPs and a carry tray to simplify the transition of the experimental setup and NHP from the procedure room to the magnet room (Fig. 1). The end-loaded dipole antenna was designed and optimized via EM simulations in XFdtd 7.5 (Remcom, State College, PA, USA) to be self-resonant at 447 MHz when loaded with a 10 cm diameter spherical phantom (relative permittivity of 56.6 and conductivity of 0.757 S/m) at a distance from the antenna of 9.5 cm (to the phantom center). The dipole was fabricated on standard FR4 PCB by a commercial board house (Advanced Circuits, Maple Grove, MN, USA). Dipoles in the 8-CH array were decoupled to −12 dB or better through careful coaxial cable management and sheath current suppression. Each dipole was matched to 50 ohms via a lattice balun^16^ at the feed point. This dipole array is used for both transmit and receive and is therefore interfaced to the system via an 8-channel (8-CH) T/R switch box which contains modular, hot-swappable T/R switch modules with integrated preamplifiers with a gain of 20 dB and noise figure of 1.0 dB. For the NHP data presented in this paper, the eight channels of this dipole array are excited with equal amplitude and CP-mode phases. For this, a single 8 kW RF power amplifier drives the array via a non-magnetic high power 8-way divider (Werlatone Inc., Patterson, NY, USA) which is located at the T/R switch box.

**Figure 1:**
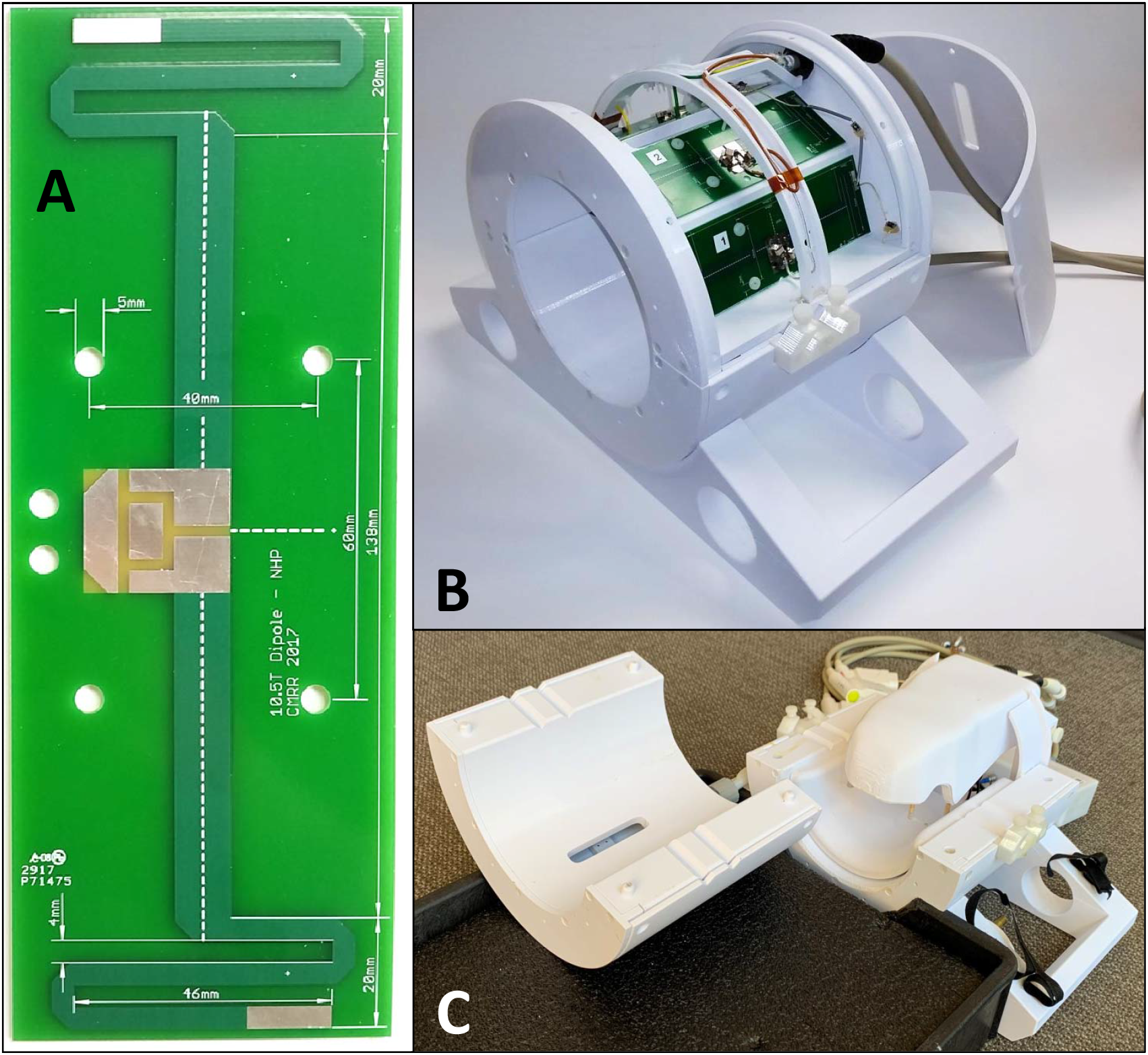
(A) Photo of the self-resonant end-loaded dipole antenna used as an element in the (B) 8-CH dipole T/R array. The top cover is removed to display the top four dipole elements. (C) The complete coil assembly with receive arrays mounted inside the dipole transceiver which in turn is mounted to a carry tray/bed. The two-halves of the splittable transceiver dipole array is shown separated from each other.

The 8-CH dipole transceiver is combined with two receive array inserts (Fig. 2): an 8-CH loop array lower cylindrical insert and 16-CH loop array constructed on a close-fitting head cap or helmet. The lower insert and the head cap coil formers were both 3D printed from PETG and have 2 mm wall thickness. The lower insert loops are rectangular 5 × 10 cm^2^ and arranged into 2 rows of 4 loops each (Figure 2 top-right) mounted on a 14.5 cm diameter cylindrical surface. The individual loops were constructed from 18 AWG (∼1.0 mm diameter) silver-coated copper wire and segmented with capacitors in six locations. Specifically, a combination of 2.7 and 3.3 pF ceramic multilayer capacitors (100B series, ATC, Huntington Station, NY, USA) and variable capacitors (SGC3 series, Sprague-Goodman, Westbury, NY, USA) located at and opposite the feed point are used for segmentation, tuning, and matching the loop. The preamps are located in an 8-CH receive box and the interconnection is made using a set of eight ¾ wavelength G_02232-09 (Huber+Suhner, Herisau, Switzerland) coaxial cables connectorized with QMA plugs (Huber+Suhner).

**Figure 2:**
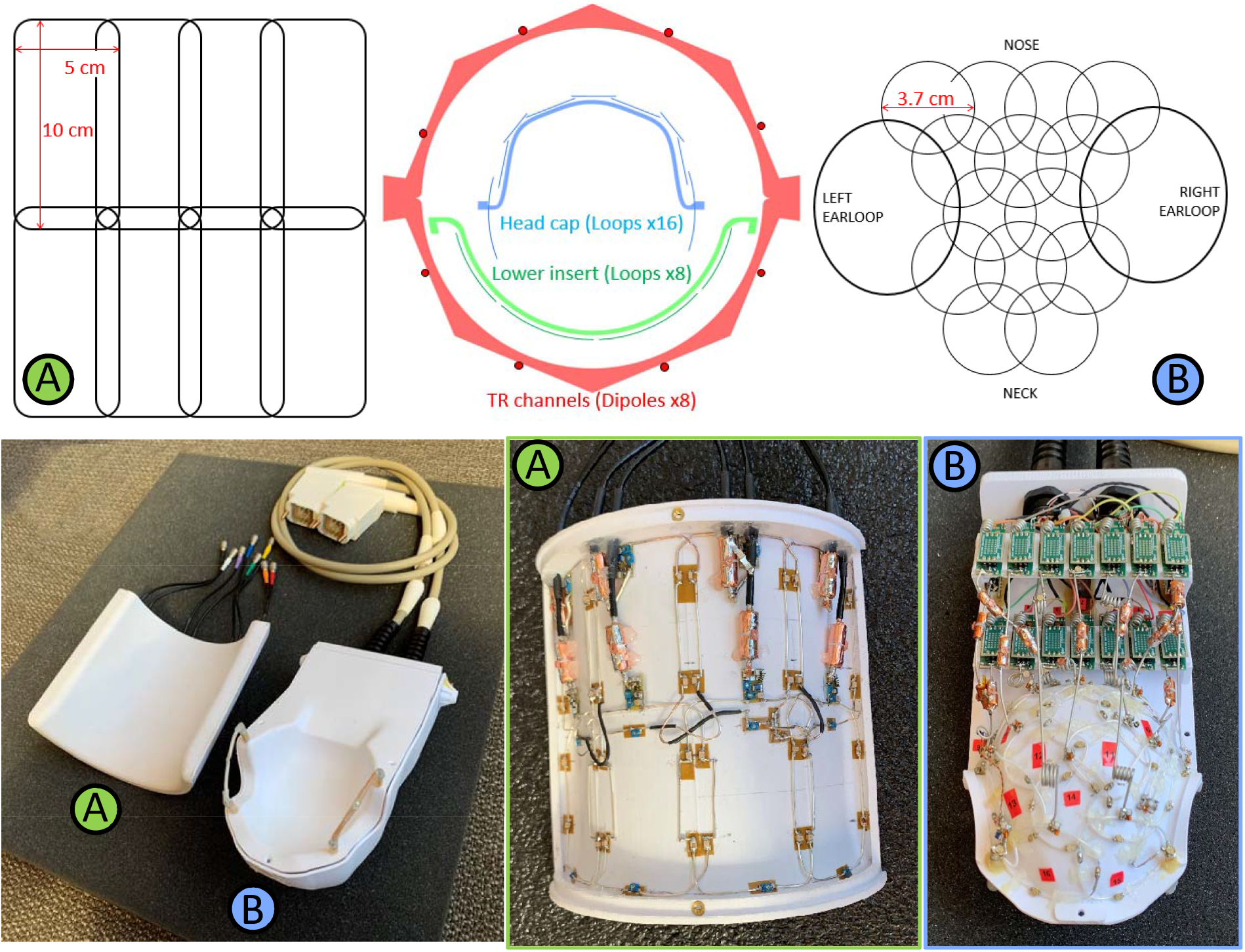
Axial cross-sectional diagram (top-center) of the coil arrangement for the dipole transmit/receive (TR array), 8-CH receive lower insert (A) and 16-CH receive headcap (B). Loop arrangements for the receive arrays are shown for the headcap (top-left) and lower insert (top-right). Also shown are housing and coils (bottom) for the lower insert (A) and headcap (B). One can also see the preamplifier boards arranged in two rows behind the head cap (bottom-right).

The conformal housing of the head cap receive array was 3D modeled based off CT scan data of an adult female rhesus macaque (Age: 10 years, weight: 6 kg). The housing has accommodations for sixteen preamplifier boards in two banks and a plate through which two cable bundles (each carrying 8 receive signals and 8 PIN diode lines) insert. Receive loops were constructed from 20 AWG (∼0.8 mm diameter) silver-coated copper wire and insulated with PTFE shrink tubing. The loops ranged from 37 to 40 mm in diameter and were arranged in rows of 4-3-2-3-2 loops with an ear loop at each end as shown in Figure 2 (top-left). Each loop has 2 to 4 segmenting capacitors with a value of 4.7 pF 0603 size SMDs (Knowles-Syfer, Norwich, UK) mounted on 0.8 mm thick FR4 PCB to provide mechanical support at these locations. A variable capacitor (9702 series Thin-Trim, Johanson Technology, Camarillo, CA, USA) is located opposite from the feed point for tuning the loop. The feed point consists of three 0603 SMD fixed capacitors, an 0807 size air-core inductor (CoilCraft, Cary, IL, USA), and PIN diode (MA4P1250NM-1072T, MACOM, Lowell, MA, USA) all mounted on 0.8mm thick FR4 PCB 5 × 5 mm^2^ in size. The full schematic is shown in Figure 3. Loops were overlapped to achieve geometric decoupling from nearest neighbors and were actively detuned during transmission to prevent harmful interactions with the transmit field.

**Figure 3:**
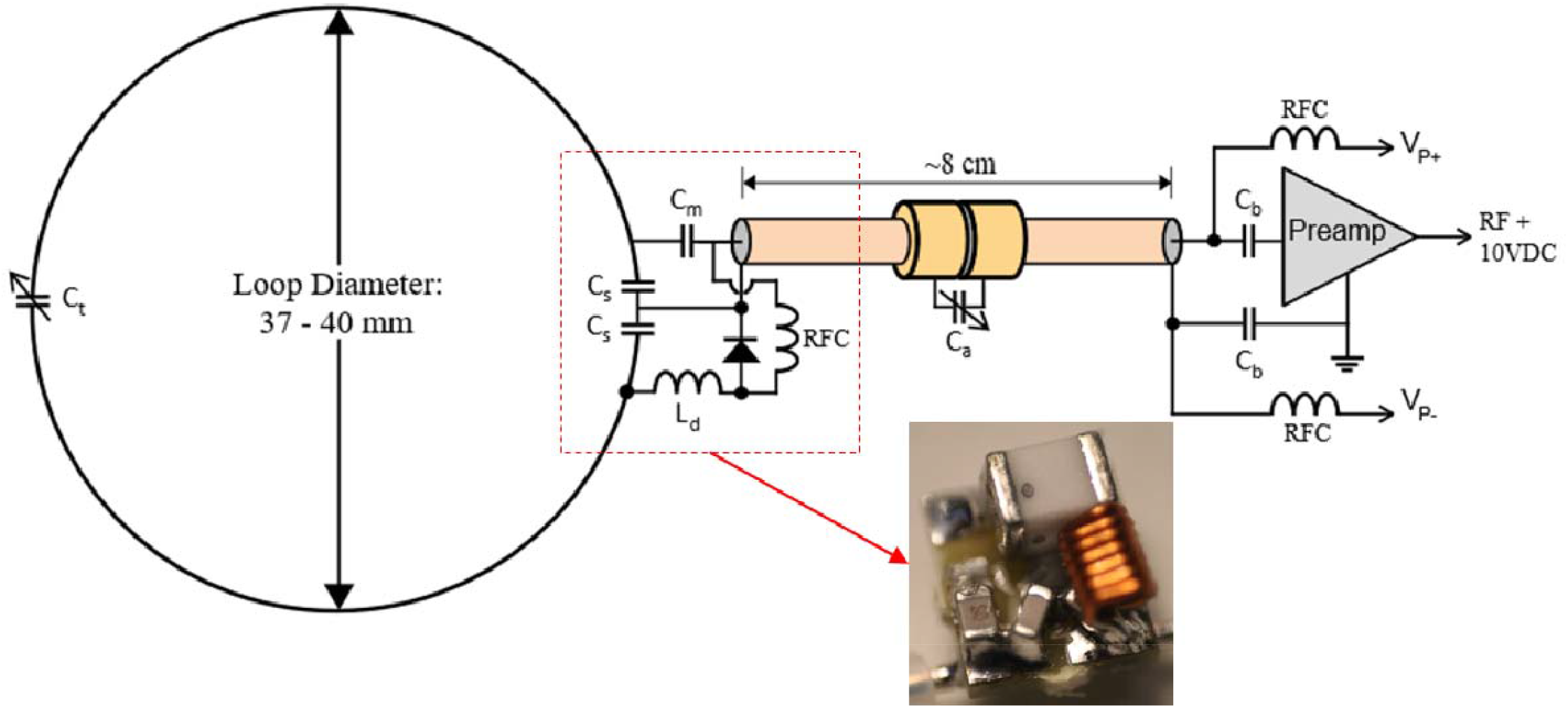
Schematic diagram of receive array channels including loop, feed point, and preamplifier. Inset: 5 × 5 mm feed board with 0603 SMT capacitors, 0807 air core inductor, and 1072T package PIN diode.

Each ear is encircled by a large receive loop which can be partially detached via a pair of non-magnetic, gold plated connectors (ODU, Mühldorf, Germany). This simplifies coil setup on an animal that has its head fixed in ear bars. Half of each ear loop is similar in construction to the rest of the 14 receive loops, however the detachable portions of the ear loops were constructed from flexible 3 mm diameter coaxial cable (only the shield is used as a conductor). Three segmenting capacitors are integrated into the detachable section and protected from mechanical stresses through a combination of the 0.8 mm thick FR4 PCB capacitor footprints and several layers of shrink tubing.

Each loop is connected to a low input impedance preamplifier via 80 mm of 1.2 mm (47 mil) diameter low loss semi-rigid coaxial cable (UT-47-C-TP-LL, Carlisle IT, USA). The length is fine-tuned for each channel to achieve the desired impedance transformation necessary for preamp decoupling. For loops located further than 80 mm from the preamp bank, slightly longer interconnecting cables are used. To compensate for the incorrect phase transformation of the longer cable, a simple phase shifter is implemented at the preamp input consisting of a single series variable capacitor. This cancels the additional phase accumulation of the slightly longer cable. For the loops furthest from the preamp bank, ¾ wavelength cables (∼24 cm) were used. These are longer than necessary to make the interconnection so excess length was coiled up.

Capacitor shortened bazooka balun cable traps^17^ were constructed along the length of each receive array coaxial cable to eliminate sheath currents that are induced by the dipole transmitter. Traps have a single trimmer capacitor across a gap located in the center of the cable trap. To tune these traps on the bench, a half-wave dipole antenna was placed in proximity and parallel to the loop coaxial cable for the channel in question and the cable trap trimmer is adjusted to minimize a through (S_21_) S-parameter measurement made from the dipole to the preamp output.

The preamplifier used is a WMA447A (WanTcom, Minneapolis, MN, USA) which was custom specified for 10.5T MRI. The preamp has 1.5 ohm input impedance, 0.45 dB noise figure, and a 28 dB gain. It is powered via the standard, system provided 10 V through the RF coaxial receive line and a bias tee. The preamp is mounted to a breakout/daughter board, which is mounted inverted to a motherboard. Both boards have an RF ground plane which the preamp is sandwiched between with the intent of providing RF shielding to the preamp from the transmit array.

The preamp motherboard is 4 × 1.3 cm^2^ in size. It contains an output cable trap and bias tee which couples the PIN diode detune signal onto the loop coax. The daughterboard and motherboard connect via a pair of 3-pin low profile headers and receptacles (BBL-103-G-E and SL-103-G-10, Samtec Inc., New Albany, IN, USA). The center pin carries the RF signal while the two endpins are RF ground.

Coil tune, detune, and coupling was evaluated on the bench using a 16-port VNA (ZNBT8, Rohde&Schwarz, Munich, Germany). The coil was then characterized on a 10.5T MRI system composed of an Agilent magnet (88 cm bore diameter, 60 cm open bore) interfaced to Siemens MAGNETOM (Erlangen, Germany) electronics. The system is fitted with whole-body gradients (Siemens SC-72) capable of 70 mT/m amplitude, 200 T/m/s slew rate, and 2^nd^ and 3^rd^ order shims (20A/channel). Noise correlation of the receive elements^18^ was calculated from a noise scan (zero RF excitation power). Parallel imaging performance was assessed as the 1/g-factor calculated for the GRAPPA technique with the quantitative GRAPPA technique^19^ using sensitivities calculated with ESPIRIT,^20^ and noise-correlation determined from noise-only data.

Ten rhesus macaque monkeys (macaca mulatta) were scanned with the proposed coil. All animal procedures were approved by the Institutional Animal Care and Use Committee of the University of Minnesota and complied with United States Public Health Service policy on the humane care and use of laboratory animals.

Monkeys receive ketamine 10mg/kg, midazolam 0.25mg/kg, and atropine 0.04mg/kg IM and were intubated and maintained on isoflurane (1-3%, inhalation) during the scan. The animal was wrapped in warm packs to maintain body temperature. A circulating water bath is used to provide additional heat. A ventilator is used to prevent atelectasis of the lungs, and to regulate CO_2_ levels. The animal is observed continuously and vital signs and depth of anesthesia are monitored and recorded at 15-min intervals.

### Acquisition parameters

T1w images and T2w images were acquired with a 3D-MPRAGE sequence and a 2D turbo spin echo sequence, respectively.

T1w: FOV: 131 × 150 mm^2^; matrix size of 280 × 320 (0.5 mm isotropic resolution); TR/TE of 3500/3.56 ms.

T2w: FOV: 112 × 150 mm^2^; matrix size of 288 × 384 (resolution of 0.4 × 0.4 × 1 mm^3^); TR/TE of 8000/68 ms; flip angle of 120 deg.

Diffusion-weighted imaging consisted of a single refocused 2D single-shot spin echo EPI sequence^21^ using a FOV of 150.8 × 83.52 × 43.5 mm^3^, matrix size of 260 × 144 × 75 (0.58 mm isotropic resolution), TR/TE of 8335/75 ms, flip angle of 90°, BW of 1860 Hz/pixel, and a phase-encoding undersampling factor of R=3 (iPAT). Diffusion-weighted images (b-value = 1500 s/mm^2^) were collected (with diffusion gradients applied along 115 uniformly distributed directions following Caruyer.^22^

Additionally, we acquired a 0.75 mm isotropic resolution for diffusion weighted images with a gradient echo sequence (Parameters: FOV: 160 × 99.68 mm^2^ (212 × 132), iPAT=3, PF=6/8, 64 slices, TR/TE = 6000/65.8ms. In 12.5 minutes, we acquire 115 diffusion directions across 2 shells (b-value of 1000 s/mm^2^ and 2000 s/mm^2^). Each of the diffusion acquisitions are repeated (FH/HF) for the purposes of eddy current and geometric distortion compensation. The entire protocol is also repeated for increase SNR.

## Results and Discussion

A noise correlation matrix for all 32 channels (Figure 4) demonstrates low levels of crosstalk (< 0.25) for most channels. A few channels have values as high as 0.6, occurring primarily in the 16-CH head cap. As expected, SNR falls off rapidly as distance from the head cap coils increase, with greatest SNR in the scalp and muscle and up to six-fold lower SNR in deep brain structures. This area of low SNR (Figure 5D, indicated in red), motivated the development of the lower 8-CH receive insert and the use of the dipole array as a transceiver. The lower receive insert and dipole transceiver each contribute approximately ¼ of the overall SNR to the deep brain region outlined in Figure 5D - 5F. As demonstrated in previous work, loop and line elements do not couple strongly and do not produce split resonances common to traditional loop transmit / loop receive coils.^14,15,23^.

**Figure 4:**
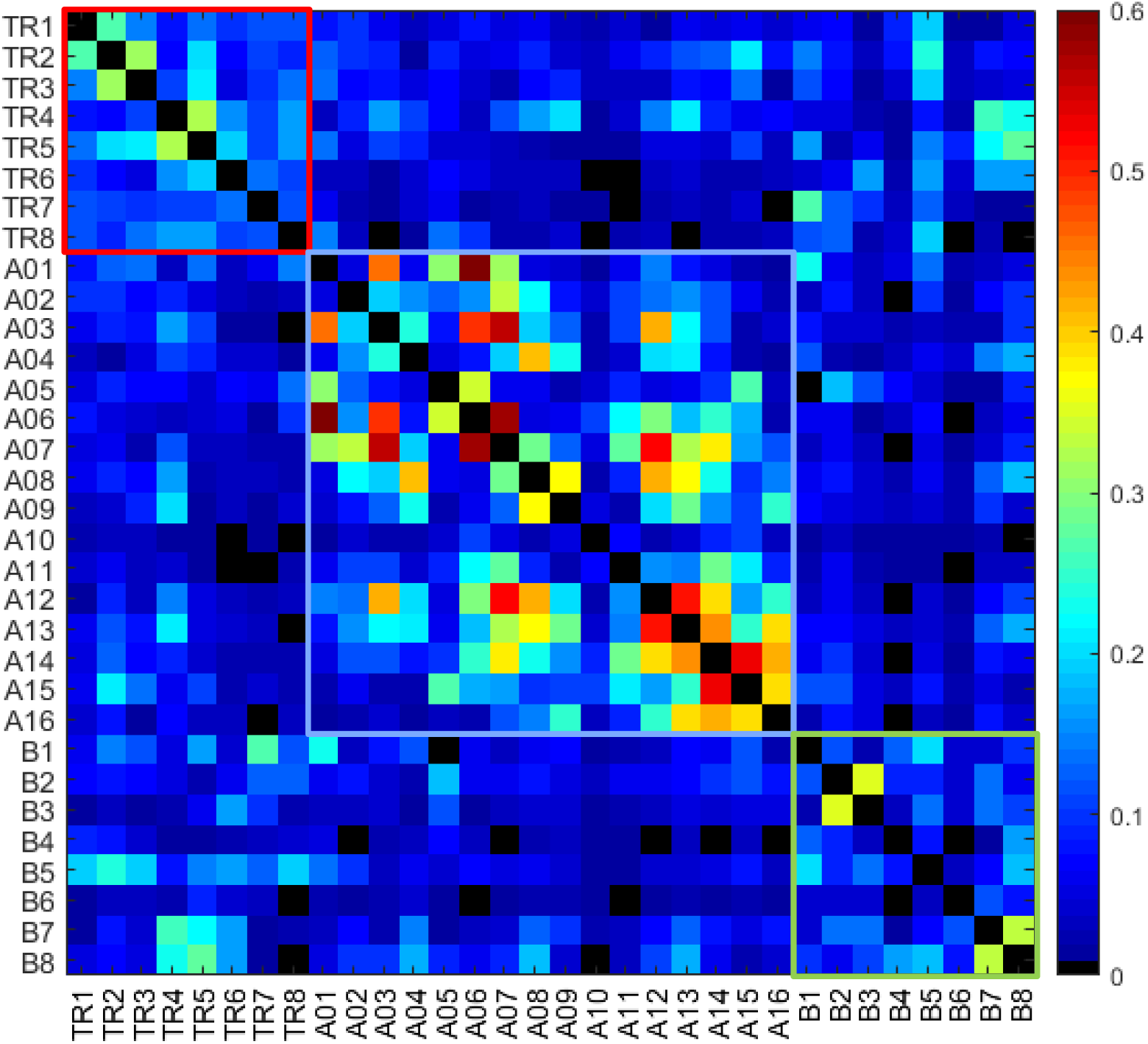
Noise correlation matrix for all 32 channels which are composed of the dipole transmit/receive channels (TR1 – 8), the 16CH headcap receiver (A01 - 16), and the lower 8CH receiver insert (B1 – 8).

**Figure 5:**
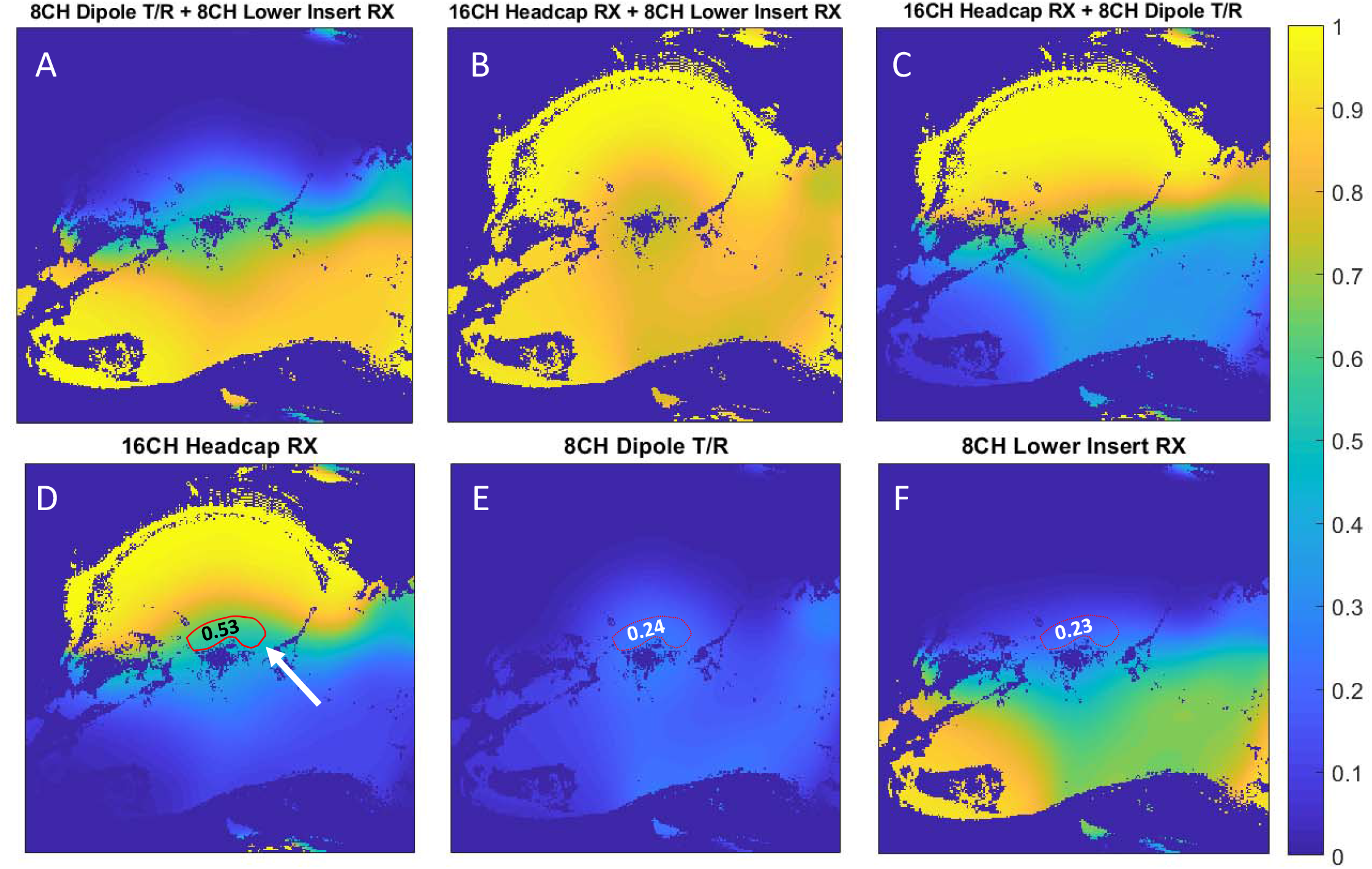
Relative SNR contribution of various combinations (A,B,C)the 16CH headcap RX (D), 8CH dipole T/R (E), and 8CH lower insert RX (F). Relative SNR contributions to the ROI drawn in red: 0.53 (16CH heapcap RX), 0.24 (8CH dipole T/R), and 0.23 (8CH lower RX).

Figure 6 shows an example of whole brain parallel imaging performance, as 1/g, for the SE-EPI with R=3, with a mean g-factor of 1.14±0.1. The inset in Figure 6, shows the histogram of the parallel imaging performance in the brain, with a maximal peak at 0.89, illustrating that on average 89% of the SNR can be retained.

**Figure 6.**
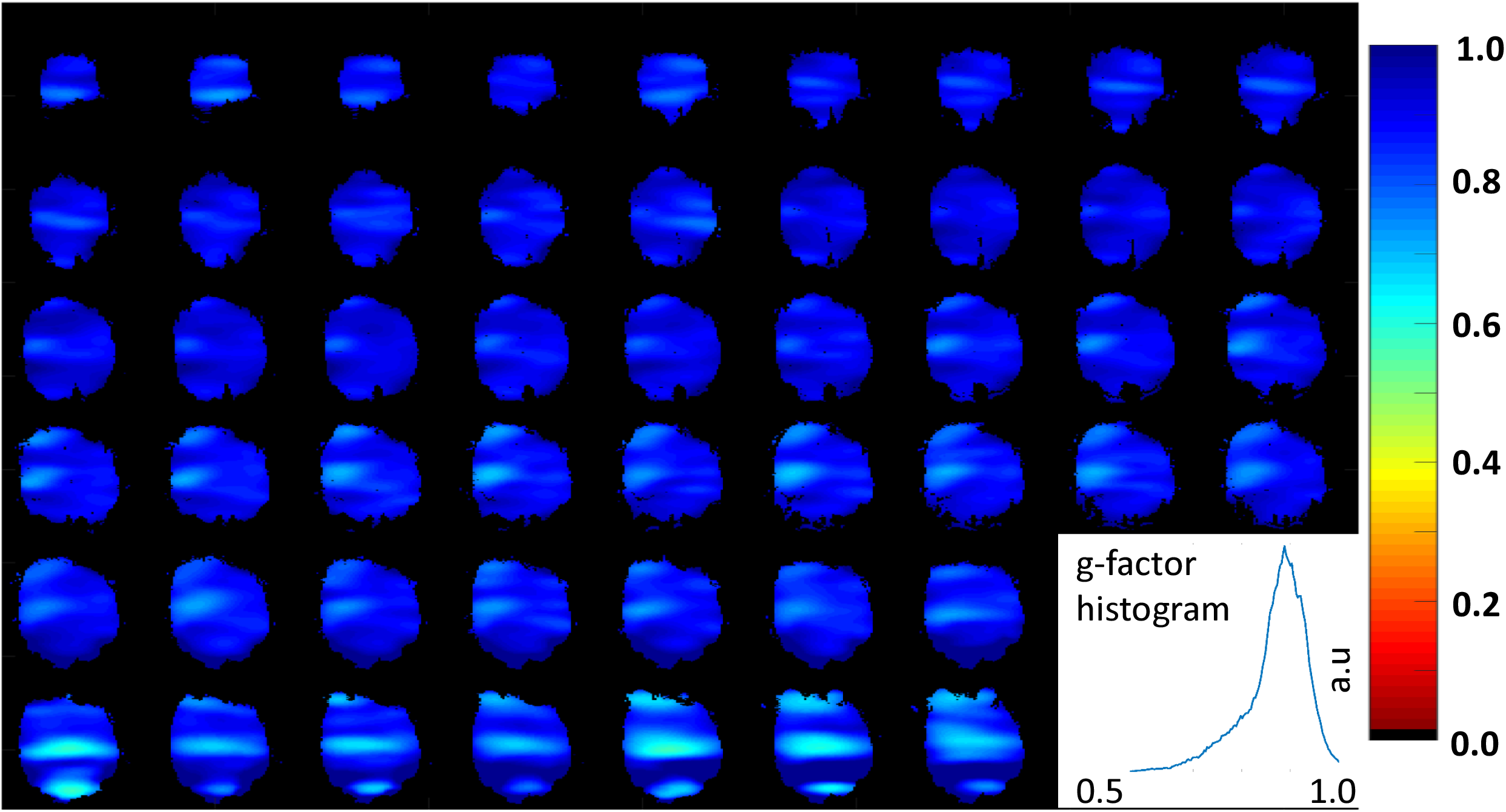
Whole brain 1/g map for SE-EPI with R=3 phase-encoding undersampling. The insert shows the histogram for pixels of the 1/g map, with a mean value of 0.89. The corresponding anatomical images are shown in Figure 8B. The quantitative GRAPPA based g-factor calculation determines a g-factor for each pixel, and an image based signal intensity mask is used to zero the g-factor outside the brain for display purposes only.

To demonstrate the utility and performance of the coil, 10 monkeys were scanned (see Methods section for details). Figure 7 & Figure 8 show representative results from such scans. Excellent T1 and T2 imaging contrast (0.5 mm isotropic and 0.4 × 0.4 × 1.0 mm^3^, respectively) was achieved at the cortical level with a sharp delineation of the white/gray matter borders (Figure 7A & B) which was maintained at deeper brain areas such as the basal ganglia region (Figure 7C). High-resolution (0.58 mm isotropic) tractography reconstruction of white-matter bundles is shown in Figure 7D.

**Figure 7:**
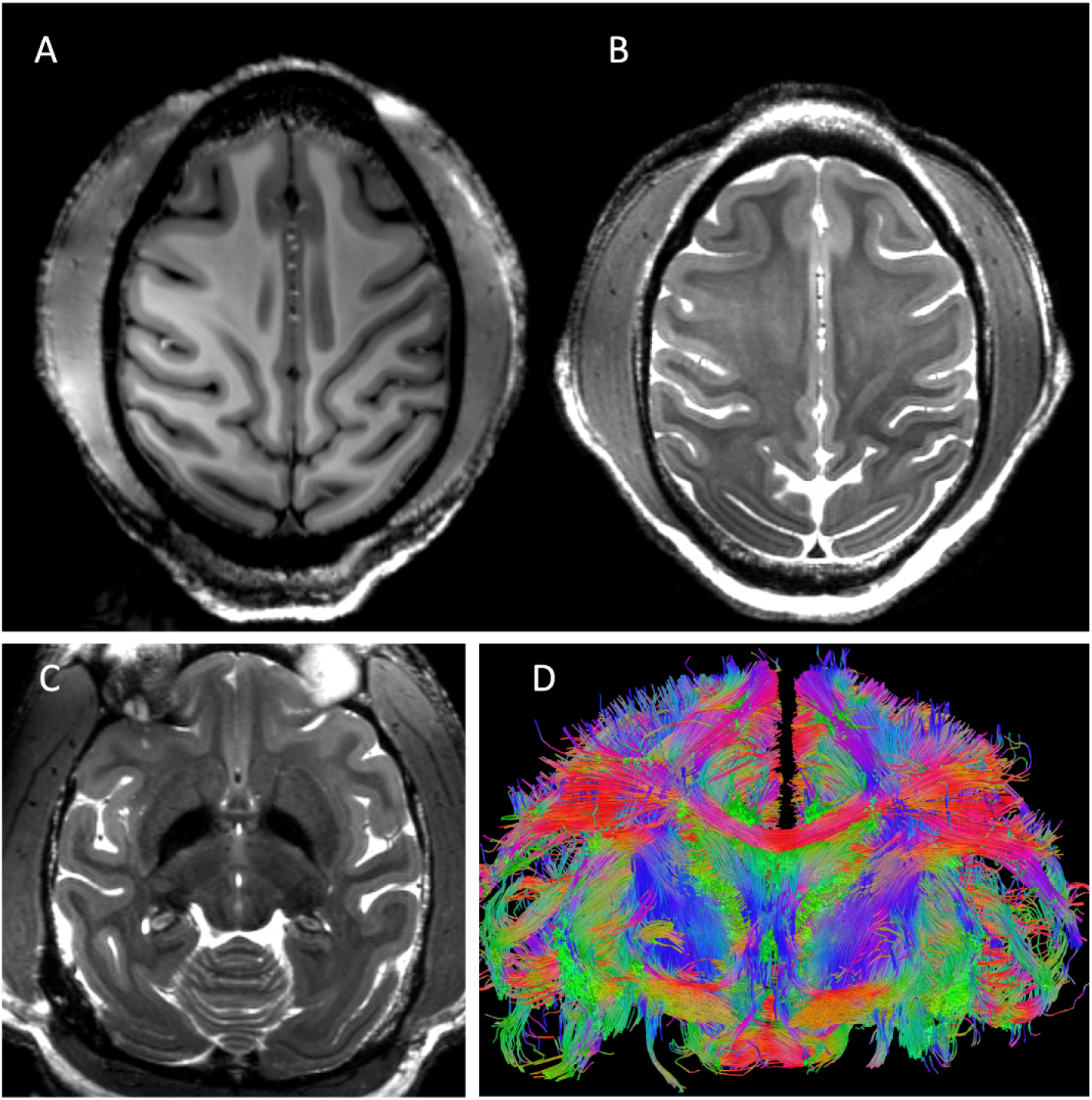
Representative results obtained with the developed coil. Excellent imaging contrast was achieved at the cortical level with (A) T1 (0.5mm iso: 131×150 mm^2^ (280×320), TR/TE=3500/3.56ms) and (B and C) T2 (0.4 × 0.4 × 1.0 mm^3^: 112×150mm^2^ (288×384),TR/TE =8000/68ms, FA=120°) weighting at 10.5T with a sharp delineation of the white/gray matter borders in superior slices (A & B) which was maintained at deeper brain areas such as the basal ganglia region (C). High-resolution tractography (0.58 mm iso) reconstruction of white-matter bundles in a coronal slice is shown in (D).

**Figure 8:**
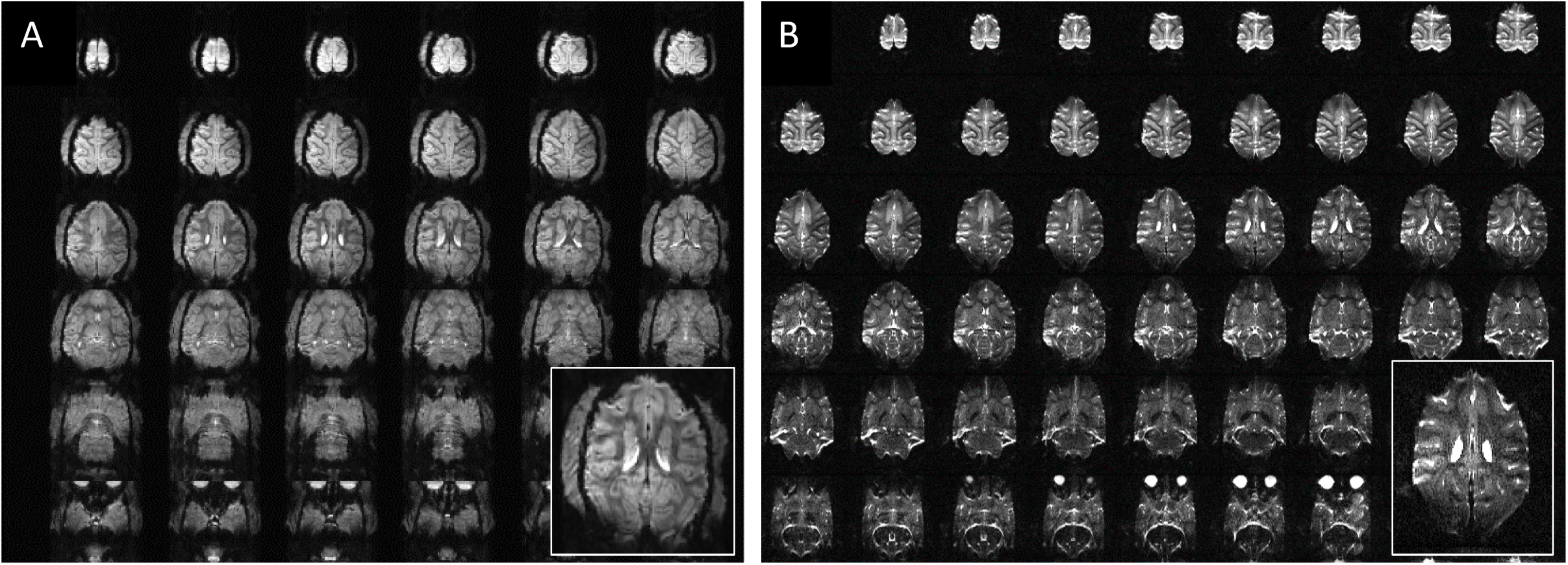
Representative examples of a high-resolution echo-planar imaging (epi) acquired at 10.5T in the NHP model. Gradient-echo (0.75 mm iso: 160×99.68 mm^2^ (212×132), TR/TE=6000/65.8ms) (A) and spin-echo diffusion (0.58 mm iso: 150.8 × 83.52 mm^2^ (260×144), TR/TE = 8335/75ms), (B) acquisition are shown.

Figure 8 shows representative examples of a high-resolution echo-planar imaging (epi) acquired at 10.5T in the NHP model, similar to the parallel imaging performance shown in Figure 6. Gradient-echo and spin-echo diffusion acquisition with 0.75 mm and 0.58 mm isotropic resolution are shown in Figure 8A & B, respectively.

## Conclusion

A 32 channel coil system for NHP brain imaging at 10.5T was developed. The coil demonstrated excellent performance and supported submillimeter high resolution imaging at 10.5T of brain anatomy, function, and tractography of fasciculated axonal bundles (obtained through, diffusion imaging (dMRI)). As demonstrated in our earlier work, the ability to combine loop and line elements was found to have numerous advantages and supported higher receive channel counts and parallel imaging gains. In particular, we demonstrated that it is possible to simultaneously receive with an array of close fitting receive loops and an eight dipole array that serves as the transmitter as well. The combined coil supported the higher acceleration factors that were essential for parallel imaging performance to support 0.6 mm^3^ high resolution fMRI and dMRI with three times reduction in sampling. Gains in SNR from inclusion of the transceive array elements during reception were more modest but nevertheless significant in the lower half of the brain and in head areas not covered by the receive loops. Future coil development will focus on further reducing the size and density of receive loops towards developing a higher channel count receive array for further gains in acceleration.

## Acknowledgements

This research is supported by NIH Grants P41 EB027061, NIH U01 EB025144, R01 NS081118, P30 NS076408 and University of Minnesota Udall center P50NS098573.

## Abbreviations

AWG: American Wire Gauge
BW: Bandwidth
CH: Channel
CNR: Contrast to Noise Ratio
CT: Computed Tomography
dMRI: Diffusion MRI
EPI: Echo Planar Imaging
fMRI: Functional MRI
FOV: Field of View
IM: Intramuscular
iPAT: Integrated Parallel Acquisition Techniques
NHP: Non-Human Primate
PCB: Printed Circuit Board
PETG: Polyethylene Terephthalate Glycol-modified
PTFE: Polytetrafluoroethylene
R: Reduction factor
RF: Radio Frequency
SE: Spin Echo
SMD: Surface Mount Device
SNR: Signal to Noise Ratio
T: Tesla
T1w: T1 weighted
T2w: T2 weighted
UHF: Ultra High Field

